# Inferring Disease Progressive Stages in Single-Cell Transcriptomics Using Weakly-Supervised Deep Learning Approach

**DOI:** 10.1101/2023.12.01.569595

**Authors:** Fabien Wehbe, Levi Adams, Samantha Yuen, Yoon-Seong Kim, Yoshiaki Tanaka

## Abstract

**Background:** Application of single-cell/nucleus genomic sequencing to patient-derived tissues offers potential solutions to delineate disease mechanisms in human. However, individual cells in patient-derived tissues are in different pathological stages, and hence such cellular variability impedes subsequent differential gene expression analyses.

**Result:** To overcome such heterogeneity issue, we present a novel deep learning approach, scIDST, that infers disease progressive levels of individual cells with weak supervision framework. The inferred disease progressive cells displayed significant differential expression of disease-relevant genes, which could not be detected by comparative analysis between patients and healthy donors. In addition, we demonstrated that pre-trained models by scIDST are applicable to multiple independent data resources, and advantageous to infer cells related to certain disease risks and comorbidities.

**Conclusion:** Taken together, scIDST offers a new strategy of single-cell sequencing analysis to identify bona fide disease-associated molecular features.

## Background

Over the past few years, single-cell technologies have rapidly advanced and been applied to patient-derived tissues to better understand and counter a variety of diseases [1]. Comparative analysis with healthy donors’ data is widely implemented to identify potential disease-associated genes [2, 3]. However, the patient-derived biospecimen is composed of a mixture of cells in various pathological stages and even contains healthy cells. Such heterogeneity obscures differential expression between patient and healthy donors and hinders identification of *bona fide* disease-associated gene expression patterns [4].

Conventionally, the cellular heterogeneity of the patient-derived single-cell data has been dissected by clustering-based approaches [5–7]. To identify disease-specific cellular states, we performed the clustering analyses (e.g. Graph-based clustering by Seurat) in single-cell RNA-seq of the midbrain from Parkinson’s disease (PD) patients and healthy young and aged donors [3]. However, uniform Manifold Approximation and Projection (UMAP) separated individual cells by cell types, irrespective of disease status (Figure S1A). The clustering divided seven cell types into 23 distinct clusters (Figure S1B), but failed to identify clusters that are unique or predominant to the patients (Figure S1C). In addition, comparative analysis between cells from PD and age-matched donors identified a limited number of up- and down-regulated genes in certain clusters (Figure S1D) and cell types (Figure S1E). In particular, the number of differentially-expressed genes was inversely correlated with the size of clusters and cell types (Figure S1D and S1F), suggesting the limitations of the clustering-based methods in the context of the classification by disease status. This statistical issue gives rise to limited differential expression of relevant PD-related genes, such as *SNCA* and *MAPT* (Figure S1F) [8]. Importantly, in PD brain, the expression of these genes shows cellular variability and is correlated with loss of neuronal connection strength [9]. Therefore, to robustly uncover disease-associated molecular elements from single-cell data, it is critical to functionally segregate cells based on disease progression levels.

Deep learning is one of the discriminative models that automatically recognize feature trends and patterns from datasets and solves a complex classification and regression problem [10]. The deep learning has been widely implemented in various single-cell data analyses, including data imputation [11], doublet identification [12], dimensional reduction [13], batch effect corrections [14], and cell type annotations [15, 16]. However, there is a limited application of the deep learning to the inference of disease progression of individual cells [17]. One of major challenges may be difficulty to train the model and regress continuous disease progression levels from the binary diagnosis information (e.g. patient=1 or healthy donor=0). To overcome such issues, we propose a novel approach, scIDST, that infers the disease progressive levels of individual cells in single-cell transcriptome profiles with weakly-supervised deep learning. The weak supervision models utilize labeling functions that are automatically generated from a small subset of labeled datasets and give weak labels on large unclear datasets [18]. We demonstrated that our weakly-supervised deep learning models successfully i) segregated cells with aberrant expression of the disease-relevant genes, ii) inferred diseased cells across different data source, and iii) inferred cells associated with comorbidity.

## Results

### Impediment of supervised deep learning in single-cell analysis

To segregate the diseased cells from the early-staged/healthy ones in PD brains, we first applied the deep learning model to the PD single-cell transcriptome data using patient information (disease diagnosis, age, and sex) as binary training data label (0 or 1) (Figure S1G) (See Supplemental Methods). Although the model separated cells of PD patients from those of healthy donors and separated cells of aged donors from those of young donors with over 93% accuracy (Figure S1H), the prediction performance for disease or age was significantly lower than that for sex (Figure S1I). Importantly, individual cells in patient-derived tissue are in various disease progressive stages and biological ages [19, 20], whereas sex is uniform among cells in one patient (Figure S1J). Therefore, the supervised binary classification model is not appropriate to infer such heterogeneous features, and it remains challenging to maximize the performance of the deep learning in the context of the single-cell data analysis.

### Weak supervision improves estimation of disease progressive levels

To overcome the conflicts between machine learning ability and single-cell analyses, we re-designed our deep learning model by employing weakly-supervised learning (Figure 1A) [21]. The weak supervision estimates the accuracy of imperfect data source and converts the binary label (0 or 1) into probabilistic label (0 ∼ 1) (See Method for detail). The probabilistic labels are calculated by Snuba/Reef system, which automatically generates the labeling functions from a small portion of the datasets and iteratively adjust parameters by pruning low-confidence labeling functions [18]. The classifier model is in turn trained from the probabilistic labels, and finally provides scores representing if a cell is likely to be diseased or aged. Unlike the supervised learning approach (Figure S1H), the weakly-supervised deep learning model outputted more broad prediction scores (Figure 1B). Particularly, the predicted disease score displayed bimodal distribution and separated cells into two groups (Figure 1C): disease progressive and early-staged/healthy cells. Importantly, vast majority of cells with higher disease score belonged to PD, and a few of them were also observed in healthy aged donors (Figure S2A). Interestingly, the high scoring cells displayed significantly-elevated expression of *alpha-synuclein* (*SNCA*), a common pathological hallmark of Parkinson’s disease (Figure 1D and S2B) [8]. In contrast, our initial supervised deep learning could not segregate *SNCA*-enriched pathological cells. Furthermore, other PD-relevant (*MAPT*, *IGF1R*, *INSR*, *HSPA1A*, *HSP90AA1*, *CNTNAP2*, and *CNTN1*) [8, 22–25], myelination-related (*MBP*, *PLP1*, and *OPALIN*), and astrocytic transporter (*SLC1A3 (GLAST)* and *SLC4A4*) genes were also differentially expressed between disease progressive and early-staged/healthy cells from PD patients (Figure 1E and S1C). In the PD brain, heat shock chaperone levels are dramatically increased in response to pathological aggregates of SNCA protein [25]. Gene Ontology (GO) analysis revealed that heat shock-related genes were significantly elevated in PD progressive cells, whereas no significant difference was observed, when globally comparing cells between PD patients and aged-matched donors (Figure 1F). Genes related to other PD hallmarks, such as neuronal death [8] and T cell activation [26], were also increased in the progressive cells (Figure 1F and S2C). In addition to disease progression, the weakly-supervised deep learning estimated biological age of individual cells. Importantly, the inferred biological age was positively correlated with *FKBP5* gene expression that is known to be elevated by aging (Figure S2D) [27]. Overall, the weakly-supervised model offers a robust platform to dissect patient-derived single-cell data and provides new insights into human disease mechanisms. We publicly provide this computational tool, single-cell Identification of Disease progressive STage (scIDST), to enable other scientists to apply the weakly-supervised deep learning model to single-cell data analysis (https://github.com/ytanaka-bio/scIDST).

**Figure 1.**
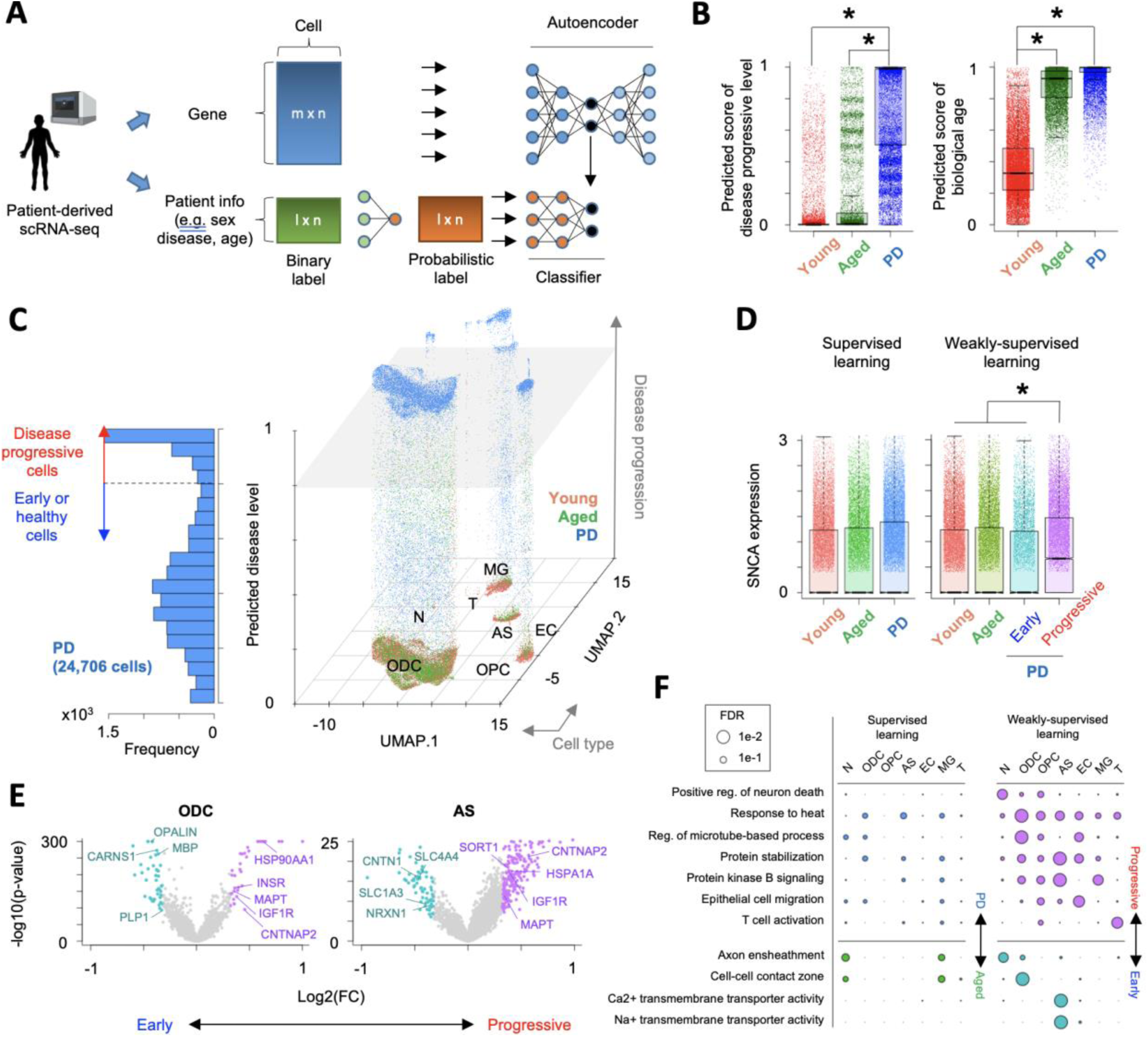
Inference of Parkinson’s disease progression levels of individual cells from patient-derived scRNA-seq. **A.** Schematic of weakly-supervised deep learning model to infer disease progressive cells. **B.** Boxplots showing predicted score of (left) disease progressive level and (right) biological age. * p<0.05 by two-sided T test **C.** 3D plot showing disease progressive cells in each cell type. X and Y-axis represent 1^st^ and 2^nd^ dimension of UMAP. Z-axis is predicted disease progressive level. Histogram of predicted disease level in PD patient-derived cells is shown in left panel. Dashed line and gray plane represent threshold of disease progressive and early-staged/healthy cells. **D.** Comparison of alpha-synuclein (SNCA) expression across groups that are segregated by (left) supervised or (right) weakly-supervised deep learning. **E.** Volcano plots showing differential expression between PD progressive and early-staged/healthy cells. **F.** Circle plot showing overrepresented GO terms (right) between PD progressive and early-staged/healthy cells, but not (left) between PD and aged brain. Circle size represents –log10(FDR). N: neuron, ODC: oligodendrocyte, OPC: oligodendrocyte precursor cell, AS: astrocyte, EC: endothelial cell, MG: microglia, and T: T cell.

### scIDST can infer disease progressive levels across multiple datasets

Given the potential of the weak supervision in the disease progressive level prediction, we tested whether our model is applicable across multiple independent single cell RNA-sequencing (scRNA-seq) datasets. Using the pre-trained model of the PD single-cell dataset (young and aged healthy donors and PD patients) (as used in Figure 1) [3], we inferred disease progressive levels of individual cells in another independent PD scRNA-seq dataset (aged healthy donor and PD patients) (Figure 2A) [6]. Similarly, the predicted disease progressive scores are broad, but significantly higher in PD patients than aged donors (Figure 2B). In addition, vast majority of cells in both aged donors and PD patients displayed high scores of biological age. To validate the reliability of the inferred disease score, we identified 284 and 193 genes, whose expression in oligodendrocyte were positively and negatively correlated with the disease progressive score in both PD datasets, respectively (Figure 2C). We observed a significant reduction in myelination-related genes (e.g. *MBP*, *MOG, PLP1*), hence confirming that the pathological feature of neurodegenerative diseases [28]. Furthermore, expression of PD- (e.g. *SNCA*, *MAPT*) [8] and other neurodegenerative disease-associated genes (e.g. *BCL2, CLU*) [29, 30] were significantly elevated with the inferred disease progression, but did not display significant difference by global comparison between PD patients and aged healthy donors (Figure S2E). These results suggest that our model did not overfit to a specific single-cell transcriptome dataset and can infer the disease progressive levels across multiple single-cell datasets.

**Figure 2.**
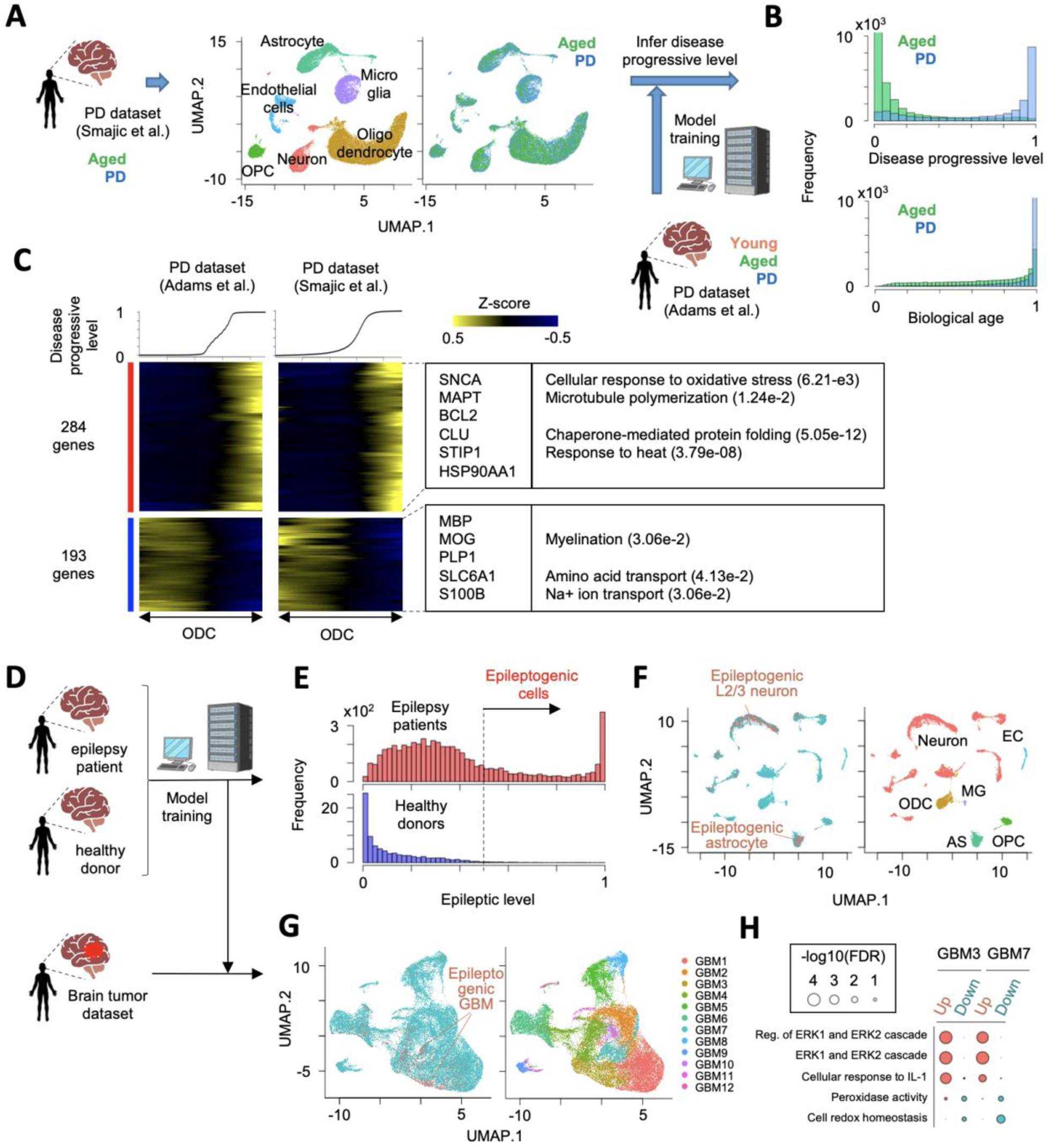
Application of pre-trained weakly-supervised models to other independent single-cell datasets. **A.** Strategy to estimate Parkinson’s disease progressive level by the pre-trained model in another independent single-cell transcriptome dataset. **B.** Histogram of predicted score of disease progressive level and biological age. **C.** Heatmap showing genes, whose expression is highly correlated with inferred disease progressive score (Genes with Pearson correlation > 0.1 in both dataset A and B are used). Oligodendrocyte cells are sorted by the disease progressive score. Heatmap color represents z-score-normalized gene expression. Representative genes and GO terms are shown in right panel. Statistical significance of GO terms is shown by false discovery rate. **D.** Strategy to infer epileptogenic cells in GBM tissues using weakly-supervised deep learning model. **E.** Histogram showing the inferred epileptic levels of individual cells in epilepsy patients and healthy donors. Dashed line represents threshold to define epileptogenic cells. **F.** UMAP plot of single cells colored by (left) epileptogenic (red) and non-epileptogenic cells (blue) and (right) cell types in scRNA-seq of epilepsy patients and healthy donors. **G.** UMAP plot of single cells colored by (left) epileptogenic (red) and non-epileptogenic cells (blue) and (right) clusters in scRNA-seq of GBM patients. **H.** Representative GO terms of differentially-expressed genes between epileptogenic and non-epileptogenic cells in GBM3 and GBM7 cluster.

To expand the practical utility of scIDST, we also tested whether our model is capable of identifying cells associated with certain symptoms and comorbidities using scRNA-seq datasets of different diseases. For example, epileptic seizure is one of the most common symptoms in glioblastoma multiforme (GBM) patients [31]. The vulnerability to seizure is positively correlated with *EGFR* level that is frequently overexpressed or amplified in GBM patients [32]. To test predictive performance of the comorbid epilepsy-related cells in GBM tissues, we first trained the weakly-supervised deep learning model with scRNA-seq of epilepsy patients and healthy donors (Figure 2D) [2]. The model clearly separated epileptogenic cells that were enriched in cortical layer 2 and 3 (L2/3) excitatory neurons and astrocytes that are major vulnerable cell types to epilepsy (Figure 2E, F, S3A, and S3B) [33, 34]. We confirmed that these segregated cells were characterized by aberrant upregulation of genes related to voltage-gated calcium ion channels, glutamatergic synaptic organization, and action potential that are hallmark of neuronal hyperexcitability in epilepsy patients (Figure S3C) [2]. Subsequently, we inferred epileptogenic cells in scRNA-seq of GBM patients using the model trained by the epilepsy datasets (Figure 2D) [35]. The pre-trained model identified the potential epileptogenic GBM cells predominantly in two clusters (GBM3 and GBM7) (Figure 2G). Differential expression analysis showed that the epileptogenic GBM cells aberrantly elevated ERK1/2 signaling genes that trigger synaptic hyperexcitation in host neural network, which in turn lead to seizure (Figure 2H) [36]. Importantly, *EGFR* is a main upstream regulator of ERK1/2 pathway. Accordingly, the inferred epileptogenic GBM cells are predominantly derived from patients with *EGFR* amplification and display significant elevation of *EGFR* gene expression (Figure S3D and S3E). Taken together, these results suggest that our weakly-supervised deep learning model has a great potential to infer cellular states associated with certain comorbidity risk and support the investigation of relationships across different diseases.

### Assessing the inferred disease progression with other pathological hallmarks

Clinically, disease progression has been often diagnosed by presence and spread of pathological biomarkers. Alzheimer’s disease (AD) is another progressive neurodegenerative disorder and characterized by amyloid beta plaques (Aβ) and neurofibrillary tangles that are primarily composed of hyper-phosphorylated Tau (pTau) protein [37]. Braak staging is one of sensitive and reliable measurements to assess cognitive and psychological status of AD patients by quantifying topographical distribution of the neurofibrillary tangles in the brain [38]. To evaluate the reliability of scIDST-based disease progression prediction, we analyzed the consistency between the inferred disease progressive levels and these clinically-used biomarkers.

We trained the weakly-supervised deep learning model with scRNA-seq of non-diseased donors and AD patients with the highest Braak stage (VI) (Figure 3A) [7]. Subsequently, the trained model was used to infer the progressive levels of AD patients with the intermediate Braak stages (I, III/IV, and V). Importantly, the inferred AD progressive levels were significantly correlated with Braak stage of AD patients (Figure 3B, Pearson correlation = 0.481, p<2.2e-16). AD-relevant genes (*MAPT* and *APOE*) were significantly elevated with the inferred AD progression (Figure 3C). In addition, average AD progressive levels in each patient were significantly correlated with quantification values of immunostaining for pTau and Aβ (Figure 3D, Pearson correlation = 0.566 (p=3.93e-3) and 0.533 (p=7.24e-3), respectively). These results indicate that the inferred disease progression by scIDST is strongly consistent with other pathological quantification methods.

**Figure 3.**
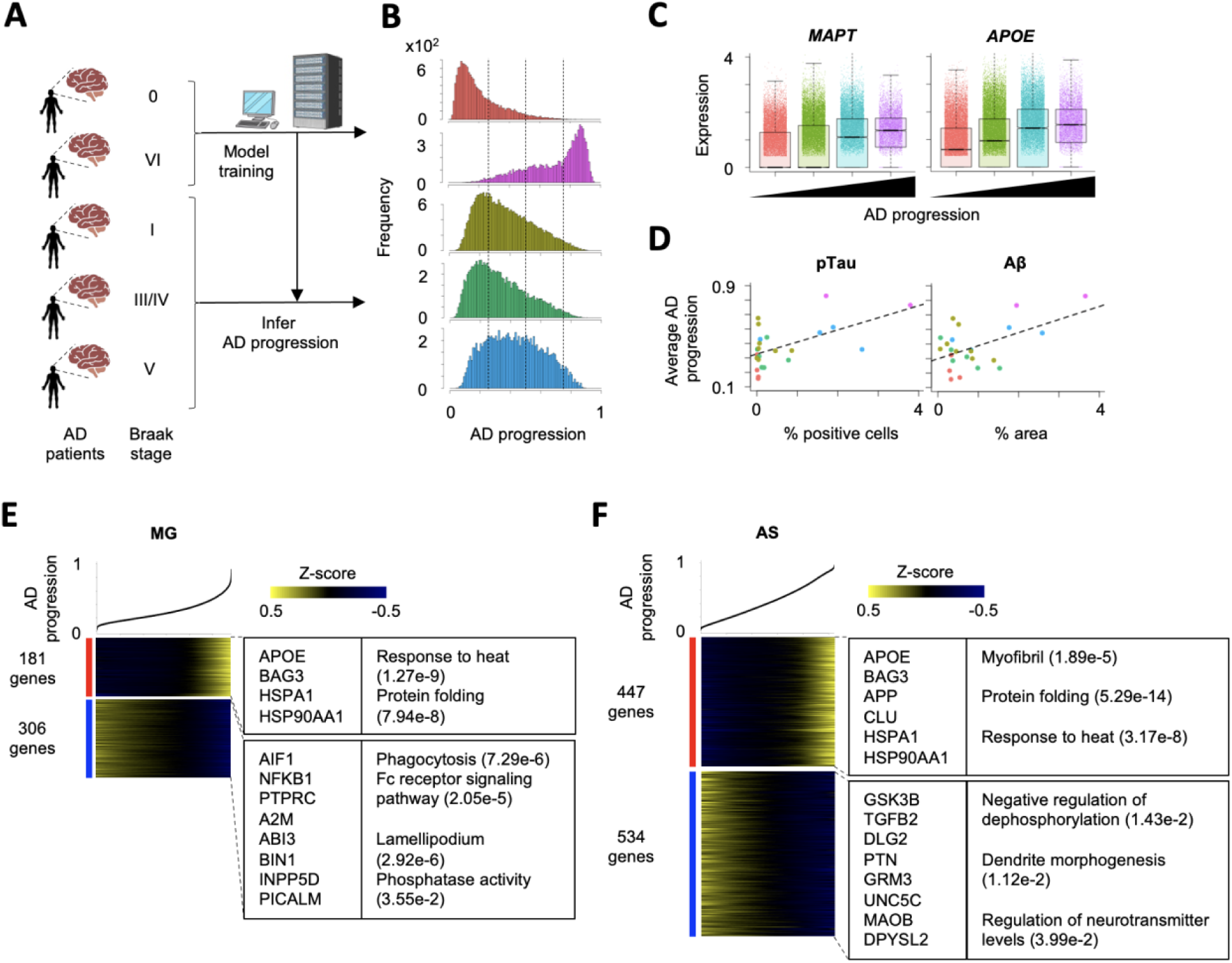
Inference of AD disease progression across different Braak stages. **A.** Strategy to estimate Alzheimer’s disease progressive levels in scRNA-seq of patient-derived samples in different Braak stages. The model was trained by Braak stage VI and 0 and estimated disease progressive levels in other Braak stages. **B.** Histograms of AD progressive levels in each Braak stage. Dashed lines are cutoff values to separate cells into four groups. **C.** Differential expression of *MAPT* and *APOE* across four disease progression groups. **D.** Comparison of average AD disease progressive levels in each patient and percentages of pathological aggregates in the brain section. **E-F.** Heatmap showing genes, whose expression is highly correlated with inferred disease progressive score (Genes with Pearson correlation > 0.1). in **(E)** microglia and **(F)** astrocyte. Cells are sorted by the disease progressive score. Heatmap color represents z-score-normalized gene expression. Representative genes and GO terms are shown in right panel. Statistical significance of GO terms is shown by false discovery rate.

AD often coincides with chronic inflammation that increases neuronal damage and is instigated by aberrant activation of microglia and astrocytes. To dissect the gene expression changes along AD disease progression, we identified 181 and 306 genes in microglia and 447 and 534 genes in astrocyte, whose expression were positively and negatively correlated with the inferred AD progression levels, respectively (Figure 3E and F). Both cell types displayed significant elevation of heat shock-related genes, which is a major characteristic of neurodegenerative diseases [25]. Interestingly, microglia significantly reduced gene expression related to phagocytosis that is essential for clearance of pTau and Aβ. In contrast, astrocytes up-regulated genes related to Aβ toxicity (*APP* and *CLU*) and down-regulated genes related to negative regulation of dephosphorylation (*GSK3B*, *TGFB2*, *DLG2*, and *PTN*) and other AD-related genes (*GRM3*, *UNC5C*, *MAOB*, and *DPYSL2*). These differential expression patterns are consistent with accumulation of pTau and Aβ in AD patients’ brains. Overall, scIDST successfully inferred AD progressive levels of individual cells, which were consistent with the pathological measurements of AD.

### scIDST also can infer cellular response to small molecules

Given the promising performance of scIDST in the inference of disease progression, we next address whether scIDST can infer other heterogeneous phenotypes, such as drug response (Figure S4) and cellular response to pathogens (Figure 4). Sexual dimorphism in brain anatomy and network connectivity emerges at developing fetal stage and is expanded with age. The relative volume of brain to body size is modestly larger in male than female [39]. The sexual bias is also relevant to the prevalence of neuropsychiatric disease onset. Females have higher risks in Alzheimer’s disease, depression and multiple sclerosis, whereas males are more likely to develop autism and schizophrenia [40]. An excellent study by Kelava and colleagues recently demonstrated that androgens specifically accelerate excitatory neuronal potential by treating an androgen steroid hormone, dihydrotestosterone (DHT), to brain organoids [41]. They also performed scRNA-seq to DHT- and mock-treated organoids, but identified only a small number of differentially-expressed genes with comparing cells between DHT- and mock-treated organoids. Furthermore, most of the differentially-expressed genes are related to ribosome biogenesis, but not associated with neuronal excitation or brain development.

**Figure 4.**
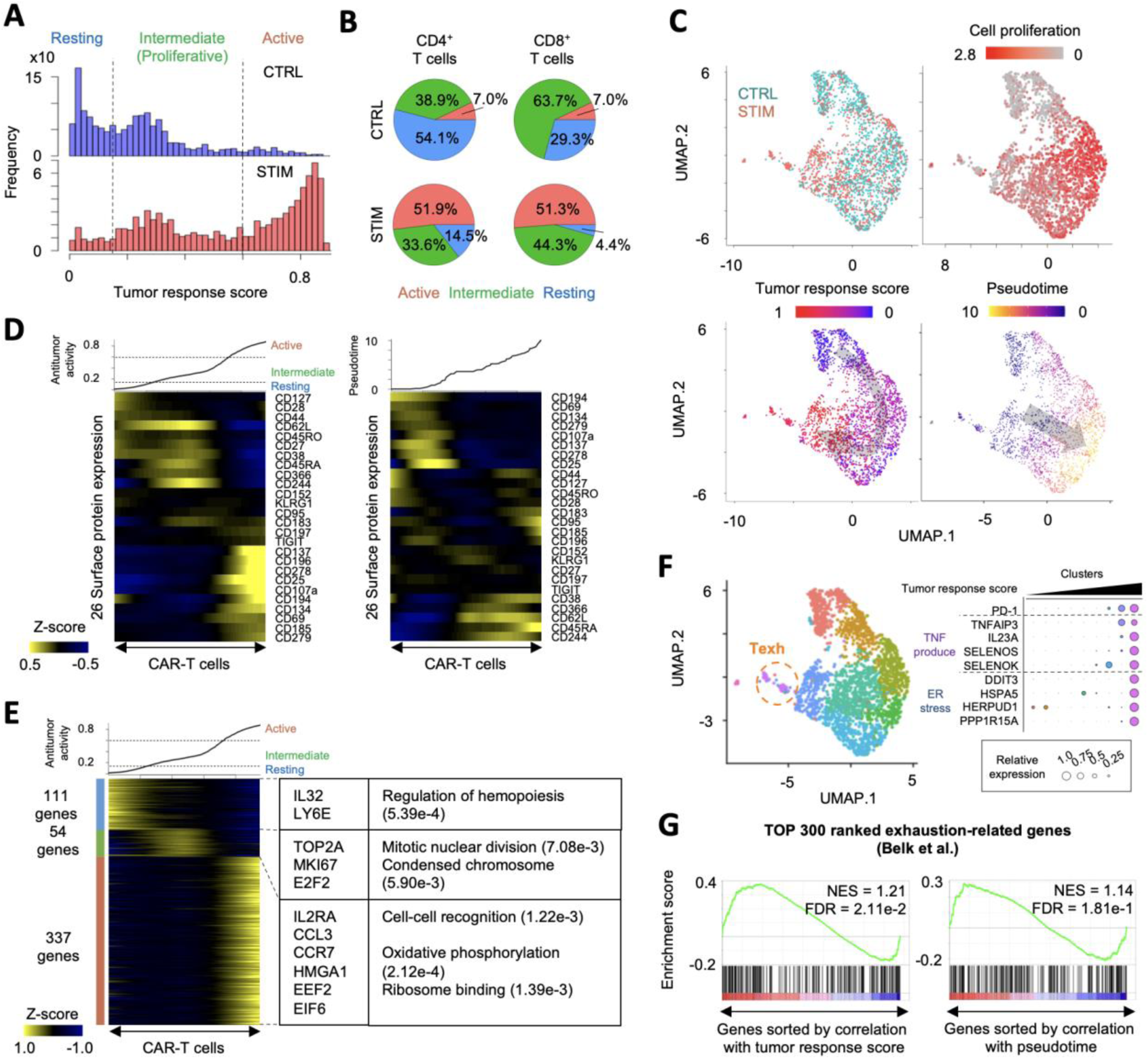
Inference of activation of CAR-T cells by neuroblastoma. **A.** Histograms showing inferred antitumor activity in neuroblastoma (STIM) and non-stimulated (CTRL) CAR-T cells. **B.** Pie chart of the ratio of three groups in CD4^+^ and CD8^+^ T cells with and without neuroblastoma stimulation. **C.** UMAP plots of individual CAR-T cells colored by the stimulation conditions (top left), cell proliferation-related gene expression (top right), the inferred antitumor activity (bottom left), and pseudotime (bottom right). **D.** Heatmap showing 26 surface protein expression dynamics across CAR T-cells. CAR-T cells were sorted by the inferred antitumor score (left) or pseudotime (right). Heatmap color represents z-score-normalized protein expression. **E.** Heatmap showing differentially-expressed genes across three groups. CAR-T cells were sorted by the inferred antitumor score. Representative genes and significant GO terms were shown in right panel. **F.** UMAP plots of individual CART-T cells colored by clusters (left). Representative differentially-expressed surface proteins and genes are shown (right). **G.** Gene Set Enrichment Analysis of T cell exhaustion gene signatures. Genes are sorted by Pearson correlation coefficient between their expression and antitumor score (left) or pseudotime (right). Normalized enrichment score (NES) and false discovery rate (FDR) are also shown.

Here, we hypothesize that only a portion of cells in the brain organoids responded to DHT, and thereby differential expression was obscured by direct comparison between DHT- and mock-treated organoids. To infer DHT response level in individual cells, we performed the weakly-supervised deep learning to scRNA-seq of DHT- and mock-treated organoids. Interestingly, a substantial number of cells in DHT-treated organoids displayed high response score (17.6% with >0.55), where cells with high score are limited in mock-treated organoids (3.99%) (Figure S4A). These DHT-responded cells were observed in specific neuronal and glial clusters (N3 and G5) (Figure S4B and S4C). In neurons, DHT-responded cells were only detected in excitatory and non-committed neurons (Figure S4D). This is consistent with Kelava *et al.*’s observation, in which inhibitory neurons were not responsible to DHT [41]. In addition, as Kelava *et al.* detected increased glial proliferation and basal progenitors in DHT-treated organoids, DHT-responded cells were detected in *TOP2A^+^* proliferating cells and *PTN^+^* basal radial glia. Importantly, these cell type-specific responses to DHT were not detectable by simple cell composition analysis between DHT- and mock-treated organoids (Left panel of Figure S4D).

Differential expression analysis between DHT- and non-responded cells identified a higher number of up- and down-regulated genes (331 and 51 genes in N3 cluster, 212 and 129 genes in G5 cluster, respectively) than global comparison between DHT- and mock-treated cells (131 and 14 genes in N3 cluster, 136 and 52 genes in G5 cluster, respectively). We detected significant differential expression of neurodevelopmental genes in DHT-responded neurons (Figure S4E). In particular, genes associated with male-biased neurological diseases (e.g. *NEUROD6* for autism, *CNIH2* for schizophrenia) [42, 43] were differentially expressed between DHT- and non-responded neurons, but not between DHT and mock-treated neurons. Overall, these results indicate that scIDST is a promising tool to identify *bona fide* differential expression in scRNA-seq analysis.

### scIDST also can infer cellular response to pathogens

Chimeric Antigen Receptor (CAR) T-cell is genetically-engineered immune cell to target specific antigens and a promising immunotherapy for cancer treatment. Clinically, the efficacy and outcome of CAR-T therapy are positively associated with expansion and persistence of CAR-T cells [44]. However, prolonged antigen stimulation exhausts T cells and leads to loss of their effector function and proliferative capacity [44]. Therefore, to overcome the limitation of current CAR-T therapy, uncovering of the molecular dynamics of CAR-T cell response is important. Here, we tested whether scIDST can infer CAR-T cell response to tumor antigens from single-cell sequencing datasets.

We first trained the weakly-supervised deep learning model with cellular indexing of transcriptomes and epitomes by sequencing (CITE-seq) of CAR T cells stimulated with or without neuroblastoma cells [45]. The inferred tumor response score displayed trimodal distribution and separated CAR T-cells into three groups: resting, intermediate, and active (Figure 4A). Both CD4^+^ and CD8^+^ CAR T-cells significantly increased the tumor response score with neuroblastoma stimulation (p<2.2e-16 by two-sided T test) (Figure 4B and 4C). Importantly, naïve or resting T cell protein markers (e.g. CD62L, CD27, CD28 and CD127) were significantly enriched in resting and intermediate group (p<1.27e-3 by two-sided T test), whereas T cell activation markers (e.g. CD137, CD25, and CD69) were significantly elevated in active group (p<5.58e-5 by two-sided T test) (Figure 4D). Differential gene expression analysis identified 111, 54, and 337 highly-expressed genes in resting, intermediate, and active groups, respectively (Figure 4E). In particular, differentially-expressed genes in intermediate group contained cell proliferation-related genes (e.g. *TOP2A*, *MKI67*, and *E2F2*). In contrast, active groups highly expressed ribosome binding genes (e.g. *EEF2* and *EIF6*) as well as genes encoding T cell activation cytokines (e.g. *IL2RA*, and *CCL3*) [46, 47]. The active proliferation followed by ribosomal activation is consistent with the dynamic changes of CAR T-cells treated in patients [48]. PD-1 (CD279) is one of major inhibitory regulators of cytokine production and effector function and leads to T cell exhaustion [44]. Since PD-1 expression was weakly elevated in active group (p<3.91e-6 by two-sided T test) (Figure 4D), we asked whether the active group also displays molecular characteristics of T cell exhaustion. Interestingly, a cell cluster with the highest PD-1 expression displayed significant up-regulation of tumor necrosis factor (TNF) production (e.g. *TNFAIP3* and *SELENOP3*) and endoplasmic reticulum (ER) stress (e.g. *DDIT3* and *HERPUD1*) that are known to contribute to the induction of T cell exhaustion (Figure 4F) [49, 50]. Furthermore, Gene Set Enrichment Analysis (GSEA) demonstrated that T cell exhaustion gene signatures were significantly elevated along the inferred tumor response score (Figure 4G) [51]. Overall, these gene and surface protein expression patterns suggest that scIDST has the capacity to infer the extent of T cell activation and exhaustion.

Cell trajectory inference is a computational method to order cells based on transcriptional similarity and characterize the molecular changes through the pseudotemporal ordering [52]. Next, we tested whether cell trajectory analysis also can infer T cell activation from the CITE-seq profiles using monocle3 [52]. The monocle3-based pseudotime value showed different patterns from scIDST-based score (Figure 4C). However, several naïve and resting markers (e.g. CD27, CD28, and CD127) was not differentially expressed through pseudotime (Supplementary Figure 5d). In contrast, pseudotime is significantly correlated with expression of cell proliferation-related genes (Pearson correlation coefficient = 0.545, p<2.2e-16) (Figure 4C). The T cell exhaustion gene signatures were also not differentially expressed along the pseudotime (Figure 4G). These results suggest that cell trajectory analysis infer the transition between proliferated and non-proliferated cells, but not T cell activation. Taken together, our results indicate that scIDST shows superior performance in predicting antitumor response of CAR T-cells.

## Discussion

Weak supervision is a new paradigm of machine learning that is optimal for large amounts of low-quality data labels, whereas other supervised learning methods (e.g. semi-supervised learning) strongly rely on the accuracy of the labels in the training datasets. Pseudotemporal ordering has been widely used to infer disease progression[3, 53]. However, it heavily relies on an inferred cell trajectory dependent on a continuous set of patient-derived samples from latent to progressive stages in order to achieve sufficient accuracy [54]. The weakly-supervised deep learning can estimate disease progressive levels without reconstruction of the cell trajectory and is capable of segregating disease-advanced cells from healthy and early-diseased stage of cells. In addition, we also demonstrated that scIDST is also applicable to infer cellular response to extracellular stimuli (e.g. drug response, immune response to cancer), suggesting the utility of scIDST for the inference of various heterogeneous phenotypes in the patients and the cultured samples.

## Conclusions

Taken together, the ability of scIDST to estimate disease status and drug response may assist in differential gene expression analysis of single-cell transcriptome data, and provide fascinating molecular insights of disease.

## Methods

### Supervised and Weakly-Supervised Deep Learning

Our weakly-supervised deep learning model is composed of three main steps: i) autoencoder-mediated dimensional reduction, ii) generation of probabilistic labels, and iii) classification of diseased cells with a multi-layered artificial neural network. The supervised deep learning model skips the probability labeling step and directly use user-given binary labels for the classifier model. Tensorflow python library (v2.9.0) with Keras Tuner API (v1.1.2) is employed to implement the deep learning [55, 56]. In the following lines, technical descriptions of scIDST are provided:

#### Preparation of single-cell data and binary data labels

As input, scIDST requires i) feature-barcode matrix and ii) binary data labels. Files of feature-barcode matrix (matrix.mtx.gz, barcodes.tsv.gz, and features.tsv.gz) can be generated by CellRanger aligner. scIDST package also provides an R script, *sgMatrix_table.R*, that can generate the feature-barcode matrix file set from Seurat object, which is a widely-used R package for quality control, normalization, and data visualization [5]. Binary data labels are manually created by users according to patient/donor information (e.g. 1 for PD patients, 0 for healthy donors) and saved as comma-separated values (.csv) format.

#### Dimensional reduction by autoencoder: autoencoder.py

The algorithm consists initially of an autoencoder that is an artificial neural network to compress scRNA-seq data into a lower dimension [57]. The autoencoder is capable of capturing non-linear relationships across data and is more appropriate for large complex datasets than other dimensionality reduction methods (e.g. PCA) [57]. Hyperparameters of the autoencoder are tunable by Keras tuner [56] to optimize the non-linear dimensionality reduction of the input: The number of hidden layers (ranging from 0 to 10 with increments of 1), the number of nodes per hidden layer (ranging from 0 to 1000 with increments of 200) and the number of nodes encompassing the latent space (ranging from 100 to 500 with increments of 100). These parameters are applied on the encoder as well as the decoder of the model. Additionally, a hyperbolic tangent activation function is employed on each layer of the autoencoder, except for the output layer of the decoder which uses a sigmoid function. The model is trained on the normalized feature-barcode matrix with 10 epochs, using an optimizer function (e.g. Adam) to minimize the mean of the sum of difference of the square between predicted output and input (i.e. the mean squared error loss function). Once the optimized weights are calculated, the encoder of the model is employed to obtain a reduced representation of the input.

A python script, *autoencoder.py*, takes the feature-barcode matrix as input and generates i) the dimensionally-reduced matrix file (.csv format) and ii) a directory storing the parameters of the autoencoder. The script also can perform the dimensionality reduction from a pre-trained autoencoder.

#### Calculation of probabilistic labels: reef_analysis.py and convert_label.py

The conversion of the binary labels to probabilistic labels is implemented by Reef/Snuba algorithm [18]. Briefly, Reef/Snuba system first generates multiple heuristics, such as decision trees, from a small portion of the dimensionally-reduced single-cell datasets and the binary labels. The confidence level of each heuristic is then calculated to prune low-quality heuristics. The Reef/Snuba performs these steps iteratively (∼50 times) and finally provides the probabilistic labels with high-quality heuristics. In scIDST pipeline, 10% of the single-cell dataset is randomly selected and used to develop the heuristics that thereafter assign the probabilistic labels to individual cells in the other 90% of the dataset. This process is repeated at multiple times (e.g. 10 times), and the average of the probabilistic labels is used for training of subsequent classifier model.

A python script, *reef_analysis.py*, takes the dimensionally-reduced data matrix (.csv format), the binary labels (.csv format) and a phenotype (e.g. disease and age), which users want to convert into probabilistic labels, as input. The output of this script is probabilistic labels of each repeat. Merging of probabilistic labels in each phenotype and the calculation of average probabilistic labels are implemented by another script, *convert_label.py*, which finally generates a matrix of probabilistic labels in csv format.

#### Classification of diseased cells: classifier_analysis.py and tensor_analysis.py

The classifier model is a multi-layered artificial neural network that can capture more complicated data patterns than single-layer network [55]. The developed sequential model is constructed with seven tunable hyperparameters. It consists of an input layer and hidden layers, which, similar to the autoencoder model, varied in i) number (2 to 10 with increments of 1) and ii) nodes containing them (50 to 500 with increments of 50). Additionally, in order to avoid overfitting, a dropout layer, with iii) variable dropout rates of 0.1 from 0 to 0.5, is introduced prior to the output layer. While a softmax is used as activation function of the output layer, iv) multiple activation functions, such as the rectified linear unit, the sigmoid and the hyperbolic tangent, are tested on the hidden layers. Finally, the model is trained with 20 epochs, using v) various optimizers (e.g. Adam, SGD) and vi) learning rates (e.g. 1e-1, 1e-2, 1e-3, 1e-4, 1e-5) to minimize viii) different loss functions (e.g. the mean squared error loss function, binary cross-entropy, categorial cross-entropy).

A python script, *classifier_analysis.py*, is composed of two submodes: *train* and *predict*. The *train* mode performs the model training on the dimensionally-reduced matrix and the probabilistic labels and gives i) predicted scores of each phenotype (.csv format) and ii) a directory of the model parameters. If “-l” option is selected, the script randomly selects a subset of data, which is used for the evaluation of the prediction performance, but not used for the model training. The *predict* mode reads the dimensionally-reduced matrix and performs the score calculation from a pre-trained classifier model.

Another python script, *tensor_analysis.py*, is an alternative command to construct the classification model, whose parameters (e.g. optimizers, loss function) can be determined by user. Similarly, the *train* mode requires the dimensionally-reduced matrix and the probabilistic labels to output the predicted score and the model directory. Since the hyperparameter tuning is skipped. This is much faster than *classifier_analysis.py*, and can be used to test the performance of weakly-supervised learning.

### Preprocessing of single-cell RNA-seq

Single-cell transcriptome data of Parkinson’s patient-derived brains and aged and healthy donors were preprocessed as described previously (GSE193688) [3]. Single-cell RNA-seq data of idiopathic Parkinson’s disease (SRP281977) [6], epilepsy brain (SRP132816) [2], brain tumor (SRP227039) [35], Alzheimer’s disease (SRP291332) [7] and dihydrotestosterone-treated brain organoid (SRP344464) [41] were downloaded from NCBI Short Read Archive (SRA). The patient information was obtained from literature or SRA meta datasheet and used to make binary labels (i.e. 0 (healthy) or 1 (disease), 0 (young) or 1 (aged), 0 (female) or 1 (male)). Raw sequence reads were aligned to human reference transcriptome (GRCh38, 2020-A) by CellRanger (v7.0.0) with default parameters. Quality control and cell clustering were then performed with Seurat (v4.0.0) in R (v4.0.5) [5]. First, we filtered out cells with 1) more than 7,000 or less than 100 detected genes, 2) more than 20,000 or less than 500 total UMI count, and 3) more than 5% of mitochondria-derived reads from subsequent deep learning analysis. Then, raw UMI count matrix was normalized to total UMI count and used to identify top 2,000 highly-variable genes that were in turn used to detect cell pairs anchoring different scRNA-seq libraries. Using these anchoring cell pairs, all scRNA-seq libraries were merged and scaled. Dimensional reduction of the merged scRNA-seq libraries was performed by principal component analysis (PCA). All cells in the merged scRNA-seq libraries were further projected into two-dimensional UMAP space using 1-30 PCs. Graph-based clustering was then implemented with shared nearest neighbor method from 1^st^ and 30^th^ PCs and 0.8 resolution value (Figure S1B and S1C). We note that the inverse correlation between cluster size and the number of differentially-expressed genes was detected, regardless of the resolution value (Figure S1D). Cell types were determined per “island” by expression of cell type-specific markers as described previously [3, 58]. Differentially-expressed genes were identified with more than 1.25-fold change and p<0.05 by two-sided unpaired T test in each cell type or cluster. Significant Gene Ontology terms in differentially-expressed genes were identified by GOstats (v2.56.0) in R packages [59].

### Assessment of binary classification models by supervised deep learning

The feature-barcode matrix of PD patients and healthy young and aged donors (GSE193688) [3] was first dimensionally reduced by *autoencoder.py.* The classifier model was then trained and assessed by the output of the autoencoder and the binary labels *classifier_analysis.py* with “-l” option. Briefly, 20% of the dataset was randomly partitioned as test dataset. The remaining dataset was further randomly divided into training and validation dataset (80% and 20%, respectively). During the training step, the model parameters were initially fit on the training dataset, and turned by the validation dataset. True positive rates and false positive rates were calculated in each category (disease, age, and sex) with test dataset at various threshold setting. Area under the curve of receiver operating characteristic curve was finally used to assess the performance of the binary classification. We repeated the above steps at 10 times and statistically compare the prediction performance across disease, age, and sex (Figure S1I).

### Application of weakly-supervised deep learning models to single-cell transcriptome profiles of Parkinson’s disease

We trained the weakly-supervised deep learning model from a single-cell gene expression of PD patients and healthy young and aged donors (GSE193688) [3]. Briefly, probabilistic labels were calculated in disease and age using the dimensionally-reduced matrix (the output of *autoencoder.py*) and the binary labels. Then, the classifier model was trained by the dimensionally-reduced matrix and the probabilistic labels. Using the trained model, we calculated scores of “disease progression” and “biological age” in two PD single-cell transcriptomic datasets (GSE193688 [3] and SRP281977 [6]). Individual cells were then separated into disease progressive and early-staged/healthy cells by 0.8 cutoff of the inferred disease progressive scores (Figure 1C). Differentially-expressed genes were identified by comparing gene expression profiles between disease progressive and early-staged/healthy cells in PD patients (Figure 1E) or age-matched donors (Figure S2C). Cells were also separated by the inferred biological ages into four groups with 0.25, 0.5, and 0.75 thresholds. Expression of a relevant age-dependent gene, *FKBP5*, was compared across these groups (Figure S2D). The correlations between the inferred disease progressive levels and gene expression were calculated by *cor* function with method=”spearman” option in R. Significant correlation was defined as more than 0.1 or less than -0.1 with p-value < 2.2e-16 (Figure 2C). GO analysis was performed in the differentially-expressed genes or the correlated genes using GOstats R package [59]. False discovery rate (FDR) was calculated by adjusting the p-value with Benjamini-Hochberg correction using *p.adjust* function with method=“BH” option in R. Less than 0.05 FDR was defined as statistical significance of GO term.

### Application of weakly-supervised deep learning models to single-cell transcriptome profiles of epilepsy and GBM patients

The weakly-supervised deep learning model was trained from a single-cell dataset of epilepsy patients and healthy donors (SRP132816) [2]. We set threshold of the inferred epileptogenic scores as 0.5, since almost all of cells from healthy donors were less than this cutoff (Figure 2E). The single-cell data of epilepsy patients and healthy donors was projected into UMAP dimension as described above (Figure 2F). Cell types were determined by cell-type specific markers. For example, L2/3 neurons were determined by co-expression of *vGLUT1* (*SLC17A7*) and *CUX2*, whereas astrocytes are by *GFAP*. Differentially expression analysis and GO analysis between epileptogenic and non-epileptogenic cells was performed in L2/3 neuron and astrocyte cluster, in which vast majority epileptogenic cells were observed (Figure S3C).

Using the trained model, we inferred epileptogenic levels in a single-cell dataset of GBM patients (SRP227039) [35]. Similarly, epileptogenic GBM cells were defined as more than 0.5 scores (Figure 2G). Enrichment of epileptogenic GBM cells were analyzed in each Seurat-based cluster. Differentially expression analysis and GO analysis between epileptogenic and non-epileptogenic GBM cells were performed in GBM3 and GBM7 clusters, in which epileptogenic GBM cells were enriched (Figure 2H and S3D). EGFR amplification status in each patient was obtained from the literature (Figure S3E) [35].

### Application of weakly-supervised deep learning models to single-cell transcriptome profiles of Alzheimer’s disease

The weakly-supervised deep learning model was trained by single-cell transcriptome profiles of Braak Stage VI (SRX9446250 and SRX9446251) and non-AD controls (SRX9446233, SRX9446234, SRX9446236, and SRX9446244) [7]. The trained model was then used to infer disease progression levels of individual cells in AD patients with Braak Stage I (SRX9446247, SRX9446248, SRX9446249, SRX9446252, SRX9446253, SRX9446254, SRX9446255, and SRX9446256), III/IV (SRX9446237, SRX9446238, SRX9446240, and SRX9446245), and V (SRX9446239, SRX9446241, SRX9446243, and SRX9446246). The percentages of pTau and Aβ-positive cells/area in each patient were obtained from the literature [7] and compared with average of the inferred disease progression scores (Figure 3D). The correlations between the inferred disease progressive levels and gene expression were calculated by *cor* function with method=”spearman” option in each cell type. Significant correlation was defined as more than 0.1 or less than -0.1 with p-value < 2.2e-16 (Figure 3E and 3F).

### Application of weakly-supervised deep learning models to single-cell transcriptome profiles of DHT-treated brain organoids

The weakly-supervised deep learning model was trained by a single-cell data of DHT- and mock-treated brain organoids (SRP344464) [41]. The threshold of cellular response to DHT was set as 0.55 (Figure S4A). After UMAP projection by Seurat, neuron (N1-5) and glia clusters (G1-11) were defined by expression of *STMN2* and *SOX2*, respectively. Neurons were further separated into excitatory (*vGLUT1* (*SLC17A7*)^+^ or *vGLUT2* (*SLC17A6*)^+^), inhibitory (*vGAT* (*SLC32A1*)^+^) and non-committed neurons (*vGLUT1^-^*, *vGLUT2^-^*, and *vGAT^-^*). Glia cells were separated into radial glia (*COL4A5^+^*), dividing radial glia (*TOP2A^+^*), and basal radial glia (*PTN^+^*) (Figure S4C). The ratio of DHT-responded cells was compared across these subtypes (Figure S4D). Differentially-expressed genes (fold change > 1.25 and p<0.05 with two-sided T test) between DHT- and non-responded cells and between DHT- and mock-treated cells (global comparison between DHT- and mock-treated organoids) was identified in N3 and G5 cluster (Figure S4E).

### Application of weakly-supervised deep learning models to CITE-seq of CAR T-cells

Gene and protein expression matrices were obtained from NCBI GEO database The count matrices of RNAs and antibody-derived tags (ADT) and CAR binder library sequencing data were downloaded from NCBI Gene Expression Omnibus (GSE181437) [45]. First, singlet, doublet, and negative cells were identified from ADT matrix with *HTODemux* function in Seurat (Doublet cells were removed from the subsequent analyses) [5]. The ADT matrix was then normalized and denoised by dsb R library (v1.0.2) [60]. Briefly, the ADT count matrix of singlet and negative cells were inputted in *DSBNormalizeProtein* function as *cell_protein_matrix* and *empty_drop_matrix* parameter, respectively. Mouse IgG antibodies were used as isotype controls. Finally, the normalized ADT matrix was combined with RNA count matrix and used for the training of the weakly-supervised deep learning models. CAR-T cells were identified by the presence of CAR binder library sequences.

The normalized RNA and ADT matrices were also used for cell trajectory analysis. Briefly, the normalized RNA data slot in Seurat object was combined with the normalized ADT matrices with *rbind* function. Subsequently, the Seurat object was converted into monocle3 object using *as.cell_data_set* function in SeuratWrappers R package (v0.3.0). Cell trajectory graph was then constructed by *learn_graph* function in monocle3 R package (v0.2.3.0) [52]. Finally, pseudotime was calculated by *order_cells* function by choosing one cluster, where the percentage of non-stimulated CAR T-cells is the highest, as root cells. The association of pseudotime with cell proliferation was analyzed by Pearson correlation between pseudotime and average expression of three major cell proliferation signatures: *TOP2A*, *MKI67*, and *E2F1* (Figure 4C)

Differentially-expressed genes were identified with more than 1.25 fold change and p<0.05 with two-sided T test by comparing one group with others (Figure 4F). T cell exhaustion gene signatures were obtained from the literature [51]. Gene Set Enrichment Analysis (GSEA, v4.1.10) of T cell exhaustion gene signatures was performed to genes sorted by Pearson correlation coefficients with the inferred antitumor score or pseudotime [61]. 0.05 FDR was used as a cutoff of statistical significance (Figure 4G).

## Availability of data and matrials

The source code and supplementary scripts are written in Python or R and publicly provided throughout our GitHub repository (https://github.com/ytanaka-bio/scIDST).

## Authors’ contributions

F.W. and Y.T. conceived the study. L.A. and Y.K. prepared single-cell transcriptome profiles of PD patients and healthy young and aged donors. L.A. performed preprocessing of PD single-cell transcriptome data. S.Y. performed preprocessing of brain tumor single-cell transcriptome data. Y.T. and Y.K. supervised the project. F.W. and Y.T. wrote the manuscript. L.A., S.Y., and Y.K. edited the manuscript.

## Acknowledgements

This research was enabled in part by support provided by Calcul Québec (https ://www.calculquebec.ca/en/) and Digital Research Alliance of Canada (https://alliancecan.ca/en).

---

**Supplemental Figure 1.**
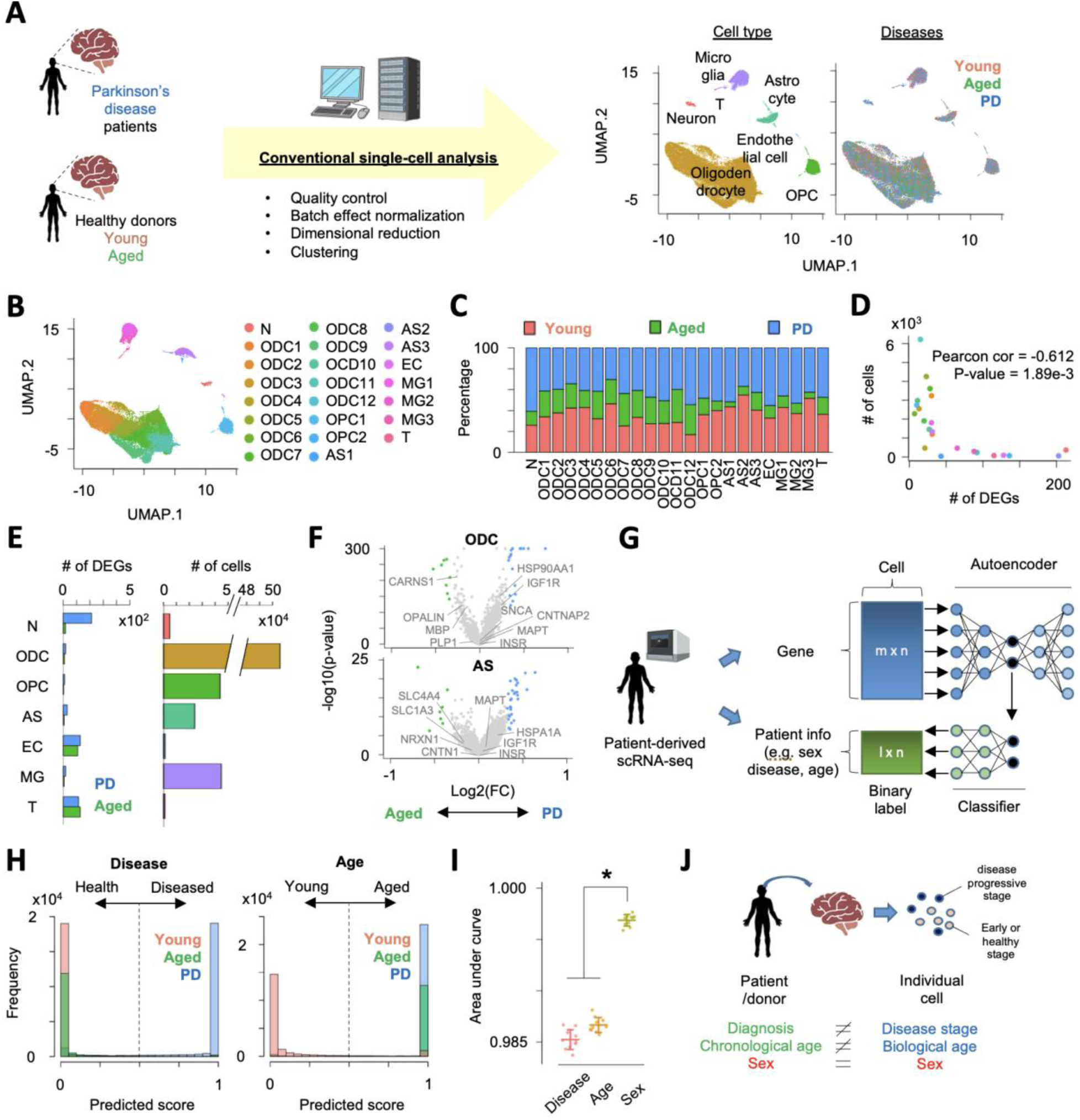
Conventional scRNA-seq analysis and application of supervised learning model to single-cell transcriptome profiles of Parkinson’s disease patients. **A.** Conventional single-cell analysis clusters cells into cell types, but not into disease**. B.** UMAP plots of single cells colored by clusters. **C.** Composition of origins of cells in each cluster. **D.** Inverse correlation between the number of differentially-expressed genes and the number of cells in each cluster. **E.** Comparison between (left) the number of differentially-expressed genes (DEGs) between Parkinson’s disease patient and age-matched healthy donor and (right) the number of cells in each cell type. **F.** Volcano plots showing differential expression between PD patients and age-matched donors **G.** Supervised deep learning model to discriminate diseased and healthy cells. **H.** Histogram of prediction score (output of deep learning model) of disease and age. **I.** Comparison of area under curve in the prediction of disease, age, and sex. Error bar represents standard deviation (n=10). * p<0.05 by two-sided T test. **J.** Difference between patient information and status of individual cells in patient-derived tissue.

**Supplemental Figure 2.**
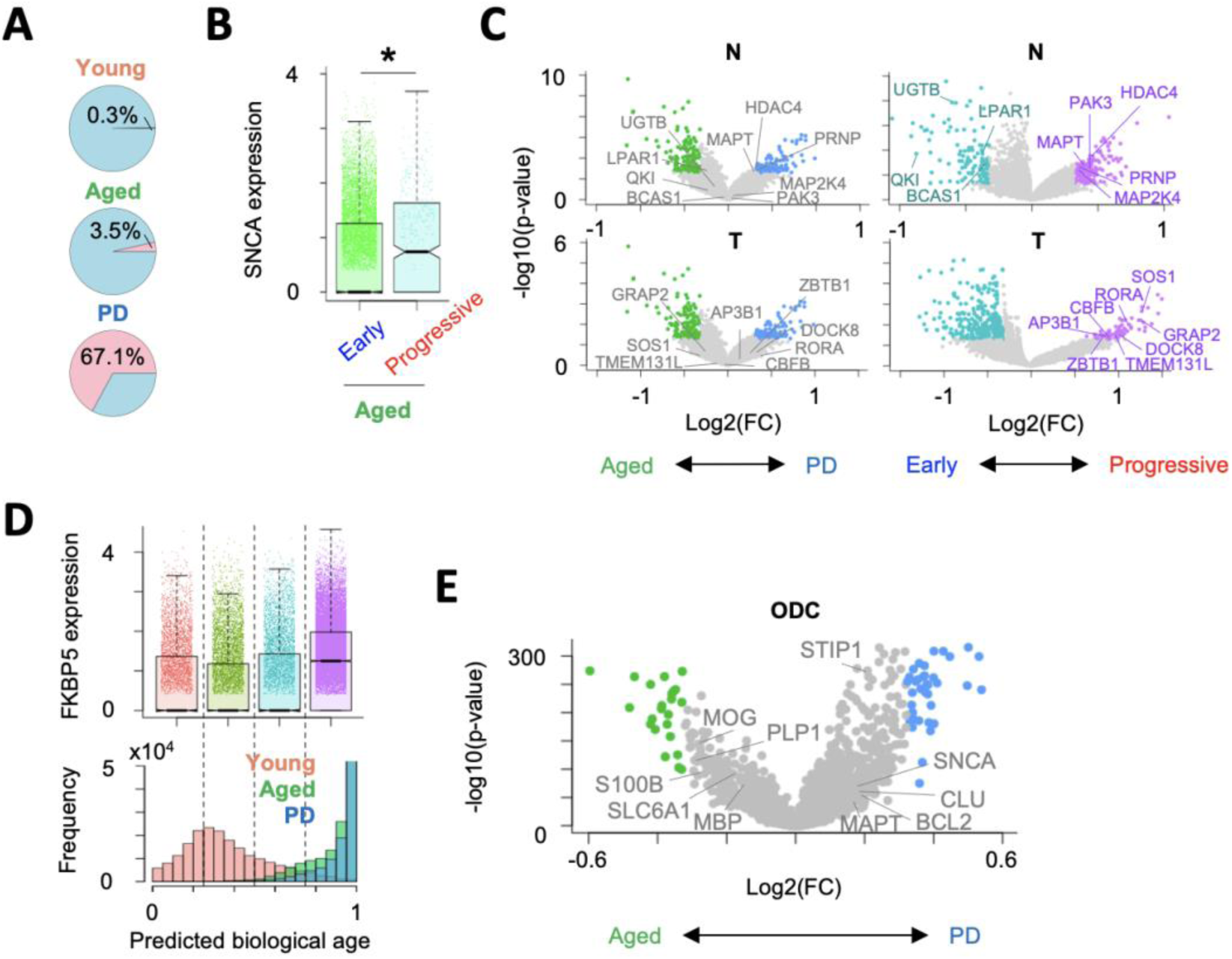
Application of weakly-supervised deep learning to Parkinson’s disease patient-derived single-cell transcriptome profiles. **A.** Pie charts showing frequency of disease progressive cells. **B.** Comparison of *SNCA* expression between early/healthy and disease progressive cells in aged brain. **C.** Volcano plot showing differential expression in neurons and T cells (left) between PD patients and healthy aged donors and (right) between disease progressive stage and early/heathy stage. **D.** Correlation between predicted biological age and *FKBP5* expression. Cells are separated into four groups by predicted biological age. Comparison of *FKBP5* expression across four groups and Histogram of predicted biological age are shown in top and bottom, respectively. **E.** Volcano plot showing differential expression in oligodendrocyte between PD patients and healthy aged donors in Smajic *et al.* dataset [6].

**Supplemental Figure 3.**
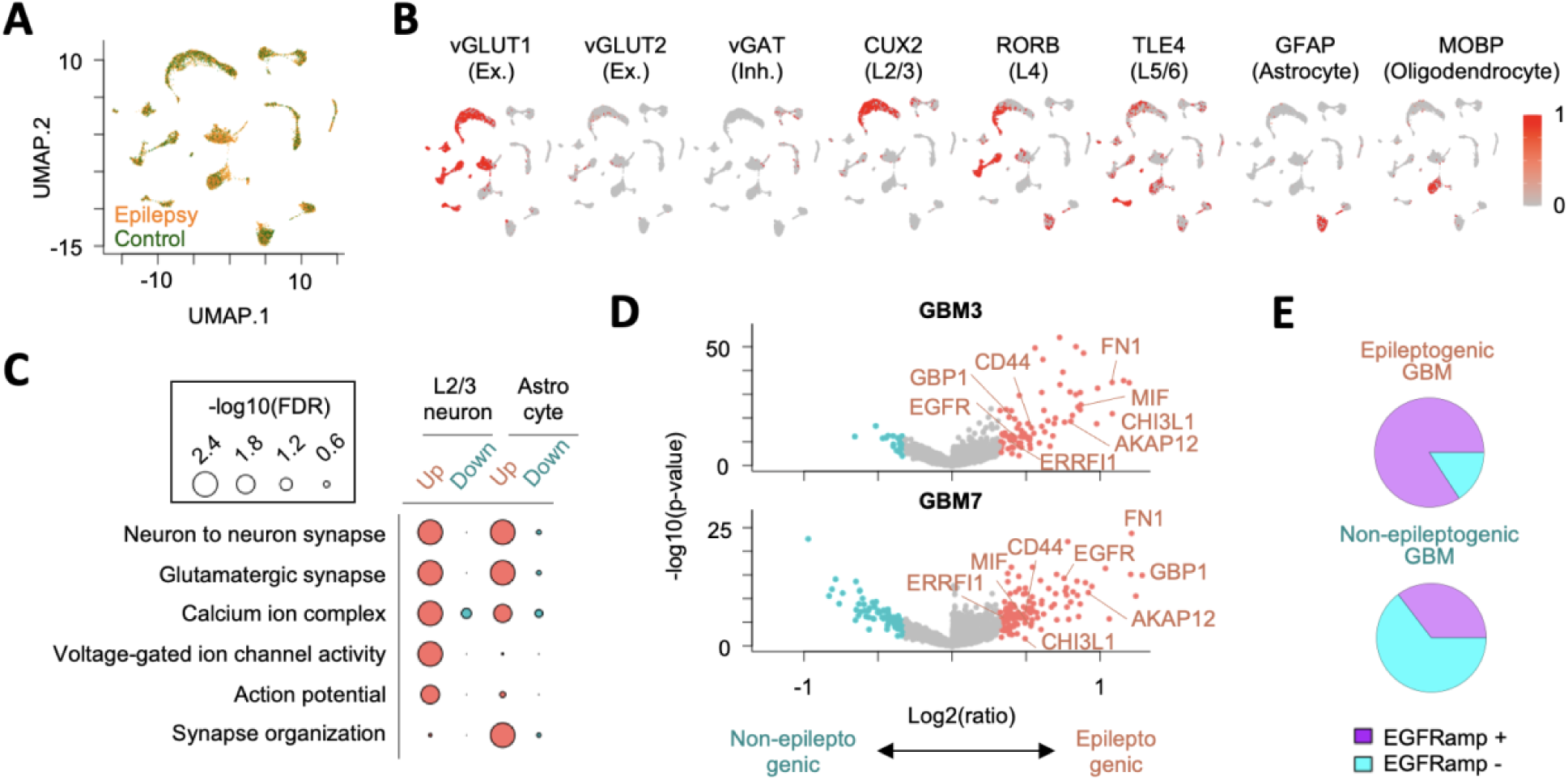
Application of pre-trained weakly-supervised models to other independent single-cell datasets. **A.** UMAP plot of single cells colored by epilepsy patients (orange) and healthy control donors (darkgreen). **B.** UMAP plot showing representative cell type-specific gene expression in epilepsy patients and healthy donors-derived scRNA-seq dataset. **C.** Representative GO terms overrepresented in differentially-expressed genes between epileptic and non-epileptic cells. Circle size represents –log10(FDR). **D.** Volcano plots showing differential expression between epileptogenic and non-epileptogenic GBM cells in GBM3 and GBM7 cluster. Significantly-upregulated ERK1/2 cascade-related genes are also shown. **E.** Pie chart showing ratio of cells with EGFR amplification in the inferred epileptogenic and non-epileptogenic GBM cells.

**Supplemental Figure 4.**
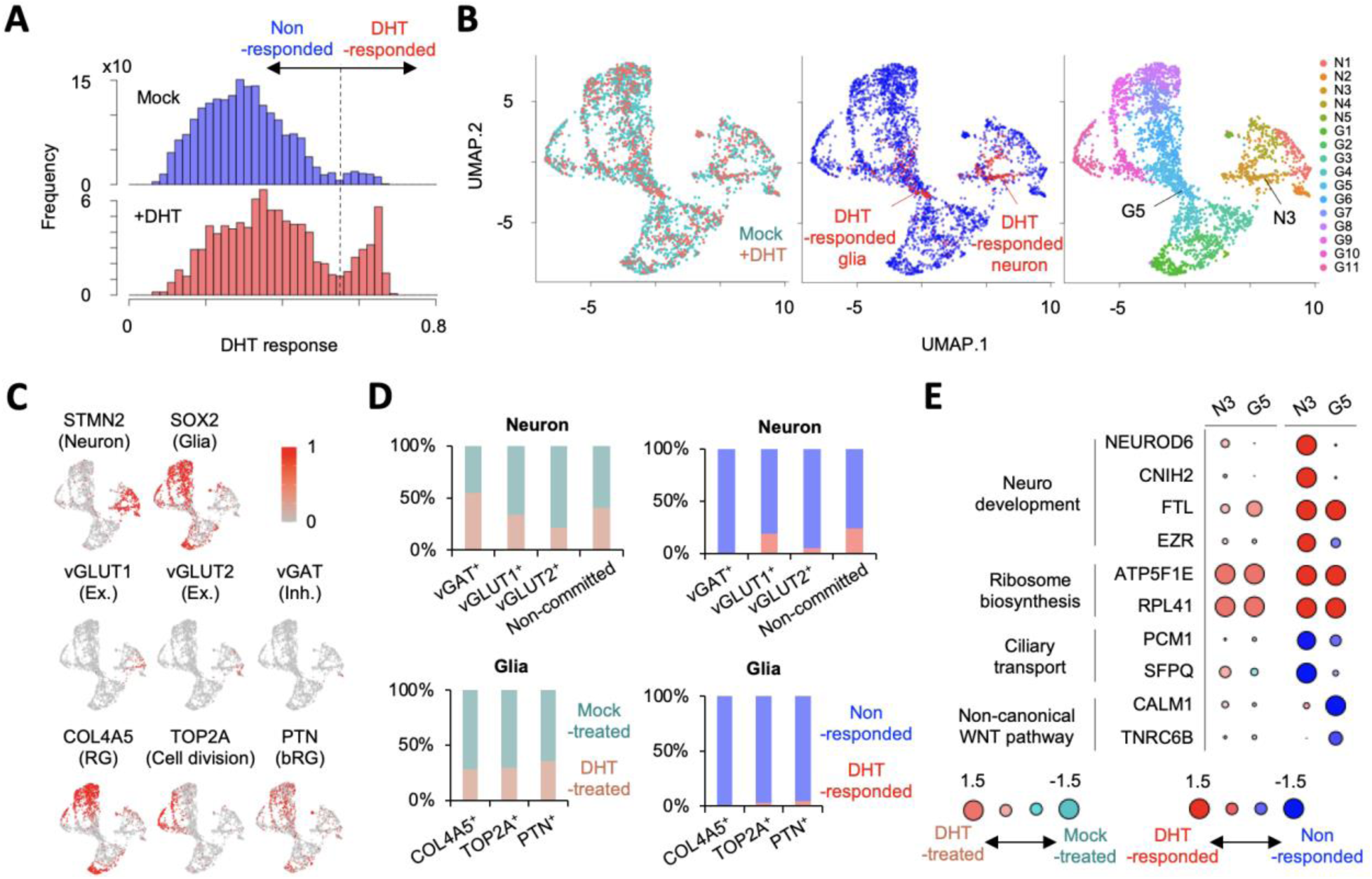
Inference of response of DHT treatment in brain organoids by weakly-supervised deep learning. **A.** Histograms of DHT response score in DHT- and mock-treated organoids. Dashed line is a threshold to define DHT-responded cells (DHT response score > 0.55). **B.** UMAP plot of single cells colored by (left) organoid type, (middle) DHT response, and (right) cluster (N: Neuron, G: Glia). **C.** UMAP plot showing representative gene expression for each cell type. **D.** Ratio of cells from (left) DHT-treated and mock-treated organoids and (right) DHT- and non-responded cells in each cell type. **E.** Representative differentially-expressed genes in N3 and G5 cluster (left) between DHT- and mock-treated cells and (right) between DHT- and non-responded cells. Circle size represent log2(ratio).

